# Altered Brain Network Topology during Successful Response Inhibition in Children with Binge Eating

**DOI:** 10.64898/2025.12.09.693025

**Authors:** Elizabeth Martin, Kurt P. Schulz, Tom Hildebrandt, Robyn Sysko, Laura Berner, Xiaobo Li

**Author notes:** Corresponding Author: Email Address, Postal Address: Xiaobo Li, PhD, 613 Fenster Hall, New Jersey Institute of Technology, New Jersey, USA.

## Abstract

**Aim:** Although a risk factor for the later development of eating disorders, few studies examine the neural underpinnings of binge eating (BE) in children. Preliminary evidence suggests a role of the corticostriatal system; the purpose of this study was to evaluate the role of the inhibitory control brain network for risk of BE in children.

**Methods:** Data from 65 children with BE (57% girls) and 84 matched controls (52% girls) from the 4.0 baseline release of the Adolescent Brain Cognitive Development (ABCD) Study were included. Stop Signal Task-based fMRI data were analyzed using graph theoretic techniques. Global and nodal network properties (e.g. efficiency, betweenness-centrality) were compared for between-group differences and sex-by-group interactions.

**Results:** Despite comparable behavioral performance, children with BE showed significantly increased nodal efficiency of the right postcentral gyrus and left middle frontal gyrus (MFG), and increased connectedness of the right postcentral gyrus compared to control children. Children with BE showed distinct network hubs including the right MFG and left insula, while controls had distinct hubs in the right orbitofrontal and left fusiform gyri. Group-specific sex differences were found in the functioning of insular and frontal cortices.

**Conclusion:** Increased efficiency and connectedness in frontal and parietal nodes of the inhibitory control network functioning in children with BE may represent a vulnerability for overeating. Distinct sex differences in functioning in children with BE compared to control children may reflect specific vulnerabilities to BE in the inhibitory control system in boys and girls that may contribute to sex differences in prevalence.

## INTRODUCTION

Binge eating disorder (BED) is the most prevalent eating disorder, characterized by discrete periods of overeating, accompanied by a loss of control over eating. In early adolescence, BED has a prevalence of 0.6-1.0%, although 6.3% of early adolescents display subclinical binge eating (BE) behaviors (1, 2). Subclinical presentations of BED in childhood and adolescence are a risk factor for the development of full-threshold eating disorder diagnoses in later life (3, 4). and extant data examining neurocognitive mechanisms underlying BE are limited. Therefore, understanding the pathophysiology underlying BE that may contribute to this trajectory is critical.

Impaired inhibitory control has been proposed as a neurocognitive mechanism contributing to BE(5). Individuals with BED and subclinical BE symptoms often have difficulty inhibiting prepotent actions when signaled (6–8), although the evidence is not consistent (9). Accordingly, disturbed function of the corticostriatal systems that subserve inhibitory control has been reported in individuals with BED (10, 11), with increased activation in widespread brain regions in adolescents with binge-type eating disorders that are distinct from both adolescent controls and adolescents with non-bingeing eating disorders (12). Further, reduced connectivity between the inhibitory control and reward networks have been reported in preadolescent children and adults with BE despite increased within-network connectivity (13, 14). The impaired inhibition of responses to rewards (e.g., food cues) presumably leads to diminished dietary restraint, which results in overeating and a perceived lack of control over consumption (15). Interestingly, deficits in inhibitory control are not limited to food-related stimuli, suggesting a broader dysfunction in inhibitory control in individuals with BE (6, 8, 16).

Alterations in neurocircuitry associated with inhibitory control may reflect an early marker of BED. Longitudinal evidence shows differences in activation during inhibitory control are evident even in children who do not have present symptoms, but go on to develop disordered eating (17). Interestingly, differences in activation are apparent during this prodromal period despite a lack of behavioral differences. It is therefore possible that dysfunction in brain regions contributing to inhibitory control may be an indicator of vulnerability to BE, even before behavioral impacts of such function are apparent.

A final possible indication that inhibitory control may contribute to BE is the existence of sex differences that emerge in prevalence of BED during adolescence and adulthood, although this difference appears less distinct prior to adolescence (18, 19). During adolescence, sex differences in inhibitory control have been widely noted (20). Given the aforementioned association of inhibitory control and BE, these sex differences may contribute to the differential risk of BE in girls and boys. Further, given that differences in brain function during inhibitory control have been noted even in the prodromal stage (17), it is possible that preadolescent boys and girls with subclinical BE may show functional differences related to inhibitory control, even if prevalence of diagnosable BED is similar.

While several existing studies have focused on either activation patterns in isolated brain regions, or connectivity between pairs of regions, inhibitory control is underpinned by a large-scale network comprising frontal, parietal and striatal regions (21). It is therefore beneficial to consider potential BE-related systems-level alterations of the inhibitory control network as a whole.

Graph theoretical techniques (GTT) allow the quantification of a functional network, providing both the ability to assess topological properties of a network as a whole, as well as the organization and contribution of individual regions (nodes) to the network (22). GTT-based analyses have begun to highlight how dysfunction in BE can be characterized by aberrance in functional network organizations (23, 24). However, despite the suggested critical role for inhibitory control in BE, and the apparent importance of the prodromal period for BE in childhood, no studies to date have investigated the systems-level characterization of inhibitory control-related functional brain organization in children with BE.

To address this gap in the literature, we compared the topological properties of the inhibitory control network in children with and without BE, using fMRI data from the Adolescent Brain and Cognitive Development (ABCD) Study baseline pool (25). Given the evidence of dysfunction in fMRI activation during inhibitory control in children and adults with BE, we hypothesized that children with BE would show distinct topological organization of the inhibitory control network during SST stop trials compared to matched control children, independent of potential behavioral differences in performance.

## METHODS AND MATERIALS

### Participants

Neuroimaging and clinical data from 90 children with BE and 112 matched controls were included in this study. These data were obtained from the Adolescent Brain Cognitive Development (ABCD) Study baseline pool (Release 4) and downloaded from the National Institute of Mental (NIMH) Health Data Archive. The ABCD Study aimed to recruit a sample that reflects the sociodemographic variation of the US population including race and ethnicity (26). The baseline pool included 11,875 children aged 9 and 10 years, from 21 sites across the United States. The ABCD study was approved by the institutional review board (IRB) of the University of California, San Diego and each data collection site. Informed consent and informed assent were obtained from parents and participants, respectively. The current study is a secondary analysis of de-identified data and therefore IRB approval was waived.

The ABCD Study baseline exclusion criteria were: a current diagnosis of schizophrenia, autism spectrum disorder (moderate, severe), mental retardation/intellectual disability, or alcohol/substance use disorder (26). We further excluded subjects with any history of traumatic brain injury or bipolar disorder to minimize confounds. Subjects with incomplete structural MRI, fMRI and/or task performance data, or low-quality imaging data (using the Human Connectome Project imaging data quality check criteria (27)) were also excluded. Based on concerns regarding design issues in the ABCD SST (28) and suggested recommendations to address these issues(29), we also excluded all subjects whose task was impacted by the noted programming error, and who had over 50% of stop trials presented with a 0 s delay.

A sample of children with BE was defined using multiple criteria. First, presence of binge eating was determined using the parent/guardian responses to the Kiddie Schedule for Affective Disorders and Schizophrenia (K-SADS) based on DSM-5 criteria (30). An initial pool of 347 children with present BE were identified, based on parent report of BE symptoms or BED diagnoses. From this initial pool, 106 subjects were excluded due to negative responses to the 7 items in the binge eating supplementary scale used to characterize binge eating behaviors (e.g. “My child eats a lot even though he or she is not hungry”, “My child feels disgusted or guilty after binge eating”). Of the remaining 241 subjects with present BE diagnosis/behaviors, we defined a group of 90 as having BE using the following inclusion criteria: i) having a BED diagnosis (n = 26); ii) having a diagnosis of BED/Other Specified Feeding or Eating Disorders/not meeting full diagnostic criteria (n = 25); or iii) no diagnosis but reported at least one binge eating episode per week for at least 3 months (diagnostically equivalent to full-threshold BED criteria) plus parent endorsement of at least one binge eating behavior on the KSADS (n = 39). **Figure 1** shows the data screening and inclusion/exclusion flow for the BE group.

**Figure 1.**
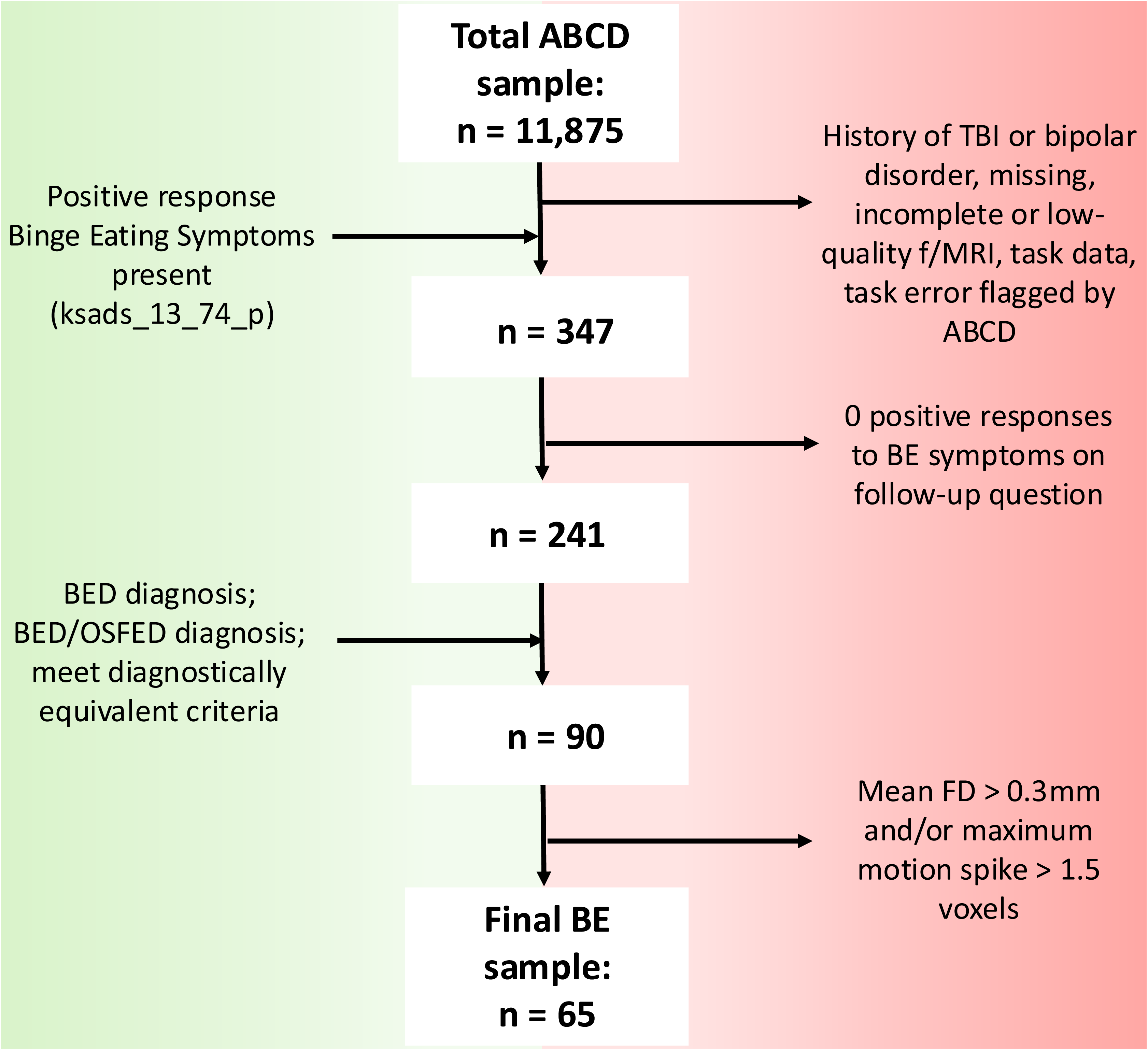
Binge eating participant inclusion/exclusion flowchart. Inclusion criteria are shown on the left (green), exclusion criteria are shown on the right (red), participants included at each stage shown in the middle (grey). BE, binge eating; BED, binge eating disorder; FD, framewise displacement; fMRI, functional magnetic resonance imaging; OSFED, otherwise specified feeding/eating disorder; TBI, traumatic brain injury.

A pool (n = 2,340) of control participants was initially identified using the exclusion criteria from the ABCD Study and the current study, who had no diagnosis of any eating disorder and no family history of psychiatric disorders. A control group of 112 participants was then defined from this pool by pseudo-randomly selecting participants matched on age, height, IQ (age-corrected picture vocabulary), sex, handedness and combined parental income (to reflect socioeconomic status) to the group with BE.

### Stop Signal Task

The SST paradigm aims to elicit the neural correlates of response inhibition. Briefly, participants respond to 180 trials of either a left- or right-pointing arrow on the screen by pressing the corresponding button (go trials). However, in 60 of these trials, the left- or right-facing arrow was replaced with an upward pointing arrow after a very short delay of < 1000 ms (stop trials). Participants were instructed to inhibit their prepotent action (i.e. inhibit the triggered press of the right or left buttons) upon seeing the upward-pointing arrow. In the ABCD sample, trials in which participants successfully withheld their response (correct stop trials), in contrast to those in which participants correctly pressed left or right (correct go) were found to elicit activation in frontal, parietal and premotor regions, as well as in caudate nucleus and putamen (31).

### fMRI Data Acquisition and Preprocessing

Data acquisition protocols from the ABCD Study have previously been published in detail elsewhere (25). Briefly, two runs of the task-based fMRI scans were acquired using whole-brain multiband echo planar imaging depicting the blood oxygenation level-dependent (BOLD) signal (TR = 800 ms, TE = 30 ms, flip angle = 30°, slices = 60, resolution = 2.4 × 2.4 × 2.4, multiband acceleration factor = 6). Raw data were downloaded from the NIMH Data Archive, data release 4.0.

Individual-level fMRI data preprocessing was carried out using the FEAT toolbox (FMRIB Software Library, FSL, version 5.0). First, the initial *n* volumes of calibration/dummy scans were removed from the time series (Philips *n* = 8, Siemens *n* = 8, GE DV25 series *n* = 5, GE DV26 series *n* = 16), to maintain 437 task-related volumes. Next, slice time correction, high-pass filtering (100 s cut-off), and head motion correction using MCFLIRT were performed on the time series. Individual task fMRI runs were excluded if framewise displacement (32) exceeded 0.3 mm and/or if maximum motion spikes exceeded 1.5 voxels. These motion-based exclusion parameters are stringent, to account for the significant effects of motion on connectivity analyses (33). Spatial smoothing using a 5mm gaussian kernel was then applied. Finally, functional images were registered to a standard space pediatric image(34), with a resolution of 2 x 2 x 2 mm.

By the end of individual-level preprocessing, a total of 53 datasets (25 from subjects with BE and 28 from the control children) were excluded from further analyses due to excessive head movements in the fMRI data.

### Subject-level fMRI Data Analysis

A model was defined for each participant in SPM12 (https://www.fl.ion.ucl.ac.uk/spm/) containing regressors for all task events: correct go trials, correct stop trials, incorrect go trials and incorrect stop trials. Twenty-four parameters were also included as nuisance regressors in the model, including the 6 basic displacement parameters (R_t_) created during rigid body realignment and the derivatives (R_t_’) and squares (R_t_ ^2^ and R ^2^, where t and t-1 refer to the current and preceding timepoints of these parameters).

### Network Node Definition

Combined activation maps including both groups were created for correct stop trials (correct stop > fixation). Network nodes were defined for each network as 4mm spheres around the peak activation in clusters of contiguous activation > 100 voxels. The correct stop network consisted of 56 network nodes including frontal, parietal, and temporal cortical nodes, and striatal, hippocampal, and thalamic subcortical nodes.

### Functional Network Construction

Beta value maps were extracted for correct stop trials, plus beta maps for the following two volumes. The total number of correct stop trials for each subject was contingent on task performance, and the number of volumes extracted was therefore different for each participant. The average number of correct stop trials across all participants was 31. The associated trial volumes were combined sequentially to form a whole-brain beta-series, and the average beta-series values for each network node were extracted. Four-level wavelet filtering was performed on the beta-series to remove motion-related noise.

Network construction and calculation of all network properties was performed using the GAT-FD toolbox in MATLAB (35). For each subject, a functional connectivity matrix (reflecting the Pearson correlation coefficient between each node pair) was first calculated. The connectivity matrix was then binarized, using a network cost thresholds such that the only connections that remain meet small world assumptions for the network (36). For a given network, cost is defined as the fraction of edges (connections between node pairs), relative to the number of all possible edges. Network and nodal properties were calculated over a range of cost thresholds from 0.1 to 0.47 in increments of 0.01, and final values used in analysis were calculated by averaging properties over calculations at all cost thresholds.

### Network and Nodal Properties

From the binarized network, the following network properties were calculated: Global efficiency (**Eq. 1**), reflecting the network’s ability to efficiently transfer information between all nodes, and local efficiency (**Eq. 2**), reflecting the network’s ability to efficiently transfer information between neighboring nodes.

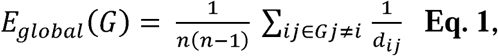

where *n* is the number of nodes, and *d_ij_* is the inverse of the shortest path length between nodes *i* and *j*.

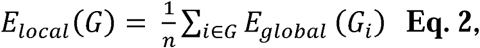

where G_i_ is a subnetwork consisting of all neighboring nodes of node *i,* and global efficiency is calculated using the equation above.

The following nodal properties were calculated for each node in the network: Nodal global efficiency (**Eq. 3**), reflecting a node’s ability to efficiently transfer information with all other nodes in a network; degree, reflecting the number of nodes any given node is directly connected to; and betweenness-centrality (BC, **Eq. 4**), reflecting the extent to which a node is necessary in the information flow between other node connections in a network. Nodes with high levels of degree or BC are considered to be hubs of a network.

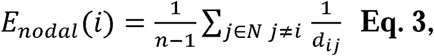

where *n* is the number of nodes, and *d_ij_* is the inverse of the shortest path length between nodes *i* and *j*.

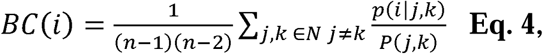

where *j, k* are pairs of nodes, *p(i|j,k)* is the probability that the shortest path between *j* and *k* passes through *i*, and *P(j,k)* is the total number of shortest paths between *j* and *k.* The steps involved in individual-level fMRI data analyses are depicted in **Figure 2**.

**Figure 2.**
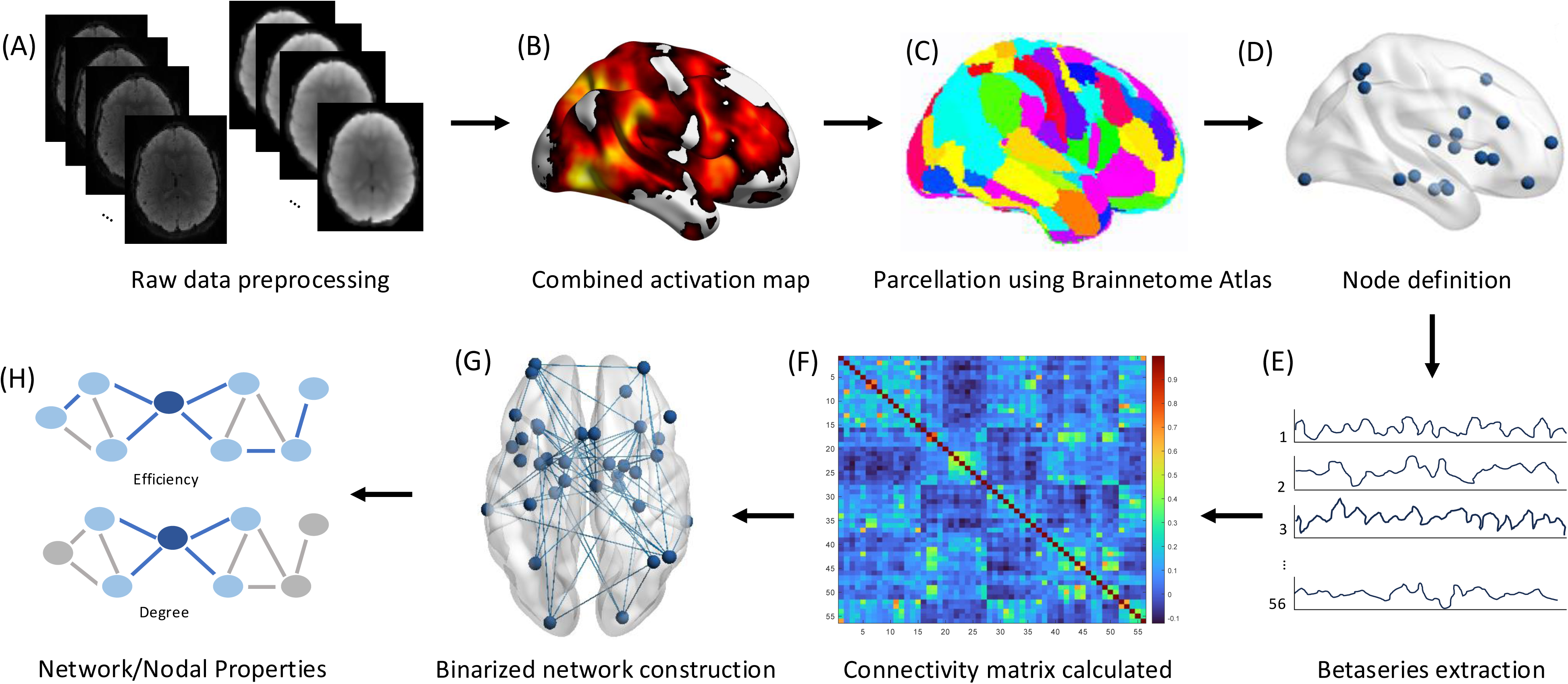
Analysis pipeline. (A) Raw data are preprocessed. (B) Group activation maps are created and combined for both groups. (C) Group activation map is parcellated using the Brainnetome Atlas. (D) Nodes are defined as 4mm spheres around independent regions containing > 100 contiguous voxels with t-values > 2.4. (E) Beta values are extracted from correct stop trials and following two volumes and combined to create beta-series, and average beta-series values for each network node is extracted. (F) Connectivity matrix is calculated for each subject and binarized. (G) The binarized connectivity matrix gives a network of nodes with or without edges between any node and all other network nodes. (H) Network and nodal properties are calculated from the binarized network of each subject.

### Group-level Statistical Analyses

Group differences in sociodemographic measures, and task performance were analyzed using an independent samples t-test. Group differences in the network and nodal properties were analyzed using the following ANCOVA model:

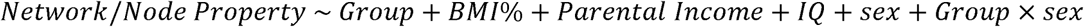

Bonferroni correction was applied for each significant node within the three nodal properties. Where nodal properties show significant group-by-sex interactions, follow-up t-tests were performed. Equivalent network properties were calculated for a go network (i.e. during performance of correct SST go trials), and are presented in **Supplementary Material**.

## RESULTS

### Demographic measures

Participants with BE were at a significantly higher BMI percentile (t(147) = 6.0, *p* < 0.001) than control participants. Participants were matched on other demographic factors. Group means and statistical comparisons of participant characteristics are shown in **Table 1**.

**Table 1.**
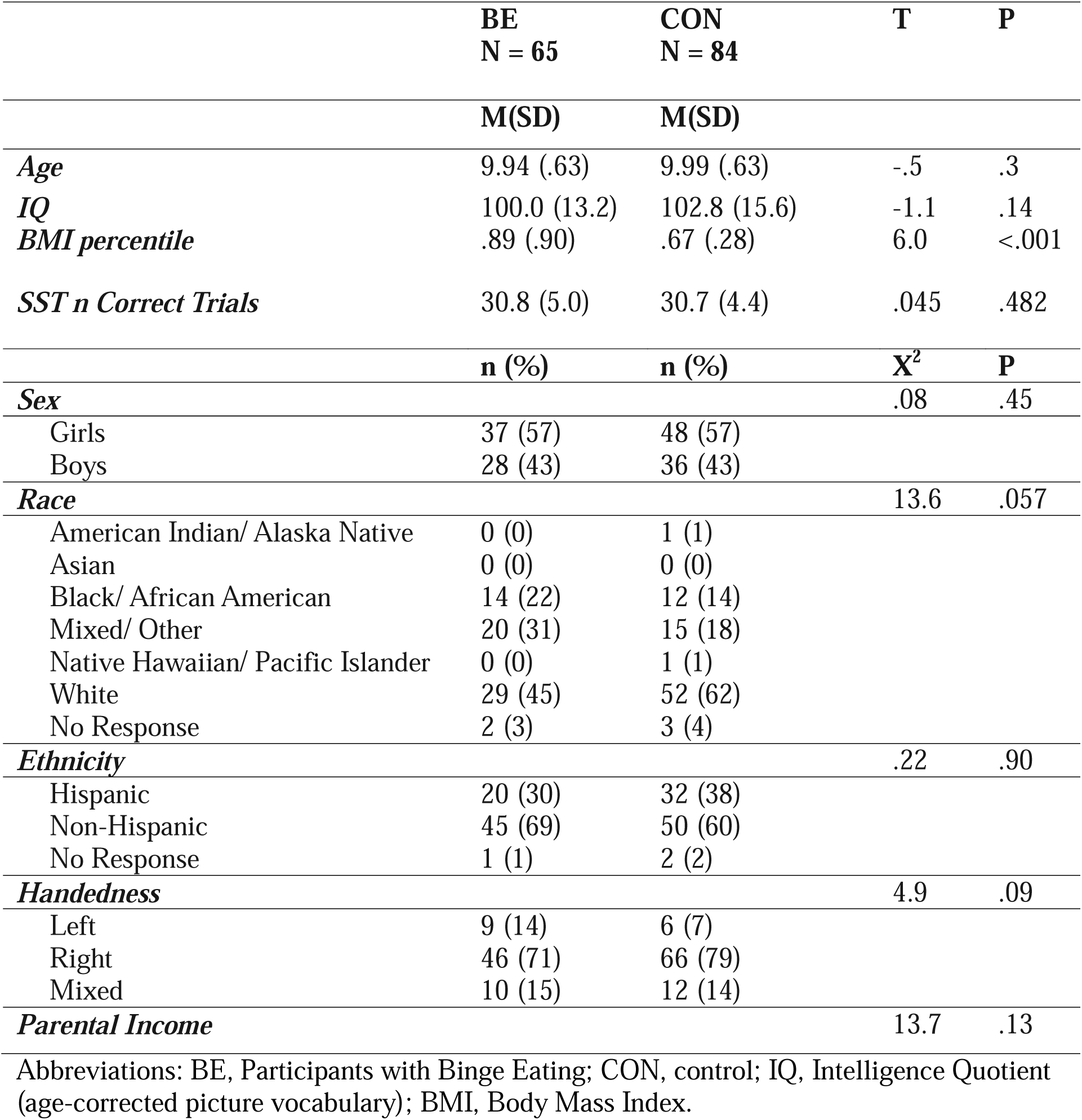
Participant characteristics and behavioral performance.

|  | **BE**<br>**N = 65** | **CON**<br>**N = 84** | **T** | **P** |
| --- | --- | --- | --- | --- |
|  | **M(SD)** | **M(SD)** |  |  |
| *Age* | 9.94 (.63) | 9.99 (.63) | -.5 | .3 |
| *IQ* | 100.0 (13.2) | 102.8 (15.6) | -1.1 | .14 |
| *BMI percentile* | .89 (.90) | .67 (.28) | 6.0 | <.001 |
| *SST n Correct Trials* | 30.8 (5.0) | 30.7 (4.4) | .045 | .482 |
|  | **n (%)** | **n (%)** | **X<sup>2</sup>** | **P** |
| *Sex* |  |  | .08 | .45 |
| Girls | 37 (57) | 48 (57) |  |  |
| Boys | 28 (43) | 36 (43) |  |  |
| *Race* |  |  | 13.6 | .057 |
| American Indian/ Alaska Native | 0 (0) | 1 (1) |  |  |
| Asian | 0 (0) | 0 (0) |  |  |
| Black/ African American | 14 (22) | 12 (14) |  |  |
| Mixed/ Other | 20 (31) | 15 (18) |  |  |
| Native Hawaiian/ Pacific Islander | 0 (0) | 1 (1) |  |  |
| White | 29 (45) | 52 (62) |  |  |
| No Response | 2 (3) | 3 (4) |  |  |
| *Ethnicity* |  |  | .22 | .90 |
| Hispanic | 20 (30) | 32 (38) |  |  |
| Non-Hispanic | 45 (69) | 50 (60) |  |  |
| No Response | 1 (1) | 2 (2) |  |  |
| *Handedness* |  |  | 4.9 | .09 |
| Left | 9 (14) | 6 (7) |  |  |
| Right | 46 (71) | 66 (79) |  |  |
| Mixed | 10 (15) | 12 (14) |  |  |
| *Parental Income* |  |  | 13.7 | .13 |

Abbreviations: BE, Participants with Binge Eating; CON, control; IQ, Intelligence Quotient (age-corrected picture vocabulary); BMI, Body Mass Index.

### Task Performance

There were no significant differences in task performance between control and BE groups. Groups means and statistical comparisons of number of correct stop responses can be found in the **Table 1**.

### Global and Nodal Topological Properties of the Inhibitory Control Network

Group comparisons showed no significant differences in the network global or local efficiencies for response inhibition (both *p* > 0.05).

Children with BE showed significantly increased nodal efficiency for successful response inhibition in the left superior frontal gyrus (SFG) (*F*(1) *=* 4.4, *p_uncorrected_ =* 0.03), left precentral gyrus (PRG) (*F*(1) = 7.4, *p_uncorrected_*= 0.007), right postcentral gyrus (*F*(1) = 10.5, *p_uncorrected_* = 0.001), right cuneus (*F*(1) = 4.9, *p_uncorrected_* = 0.028), right occipital gyrus (*F*(1) = 5.3, *p_uncorrected_* = 0.02), right thalamus (*F*(1) = 5.8, *p_uncorrected_* = 0.02), left middle frontal gyrus (MFG) (*F*(1) = 9.8, *p_uncorrected_*= 0.002), left inferior temporal gyrus (*F*(1) = 5.8, *p_uncorrected_* = 0.02), left inferior parietal gyrus (IPG) (*F*(1) = 4.4, *p_uncorrected_* = 0.04) and left insula (*F*(1) = 5.9, *p_uncorrected_*= 0.02) relative to controls. Only the group differences in right postcentral gyrus and left MFG remained significant following Bonferroni correction **(Figure 3**; **Table 2)**. BMI did not significantly contribute to the ANCOVA model for nodal efficiency in the right postcentral gyrus or left MFG (ps > 0.05).

**Figure 3.**
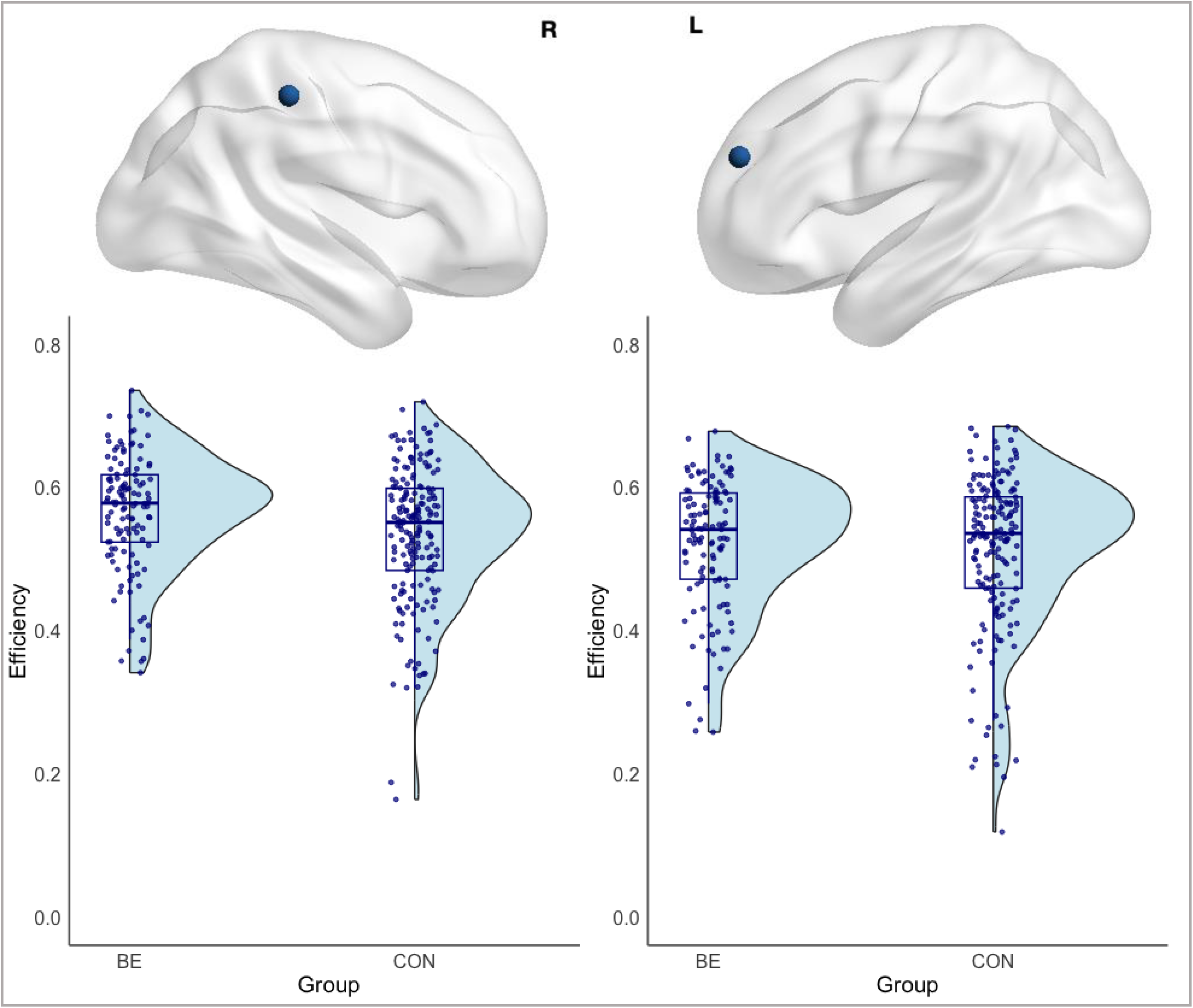
Group data for efficiency in the right postcentral gyrus (A) and left middle frontal gyrus (B). Boxplots show the median and inter-quartile ranges of the data. BE, binge eating; CON, control.

**Table 2.** Nodes showing significant (uncorrected) differences in network nodal properties.

| Region | BE<br>vs<br>CON | BE<br>M(SD) | CON<br>M(SD) | p<sub>uncorrected</sub> | p<sub>BON</sub> |
| --- | --- | --- | --- | --- | --- |
| *Nodal Efficiency* |  |  |  |  |  |
| L Superior Frontal Gyrus | ↑ | .57 (.06) | .55 (.10) | .036 | NS |
| L Precentral Gyrus | ↑ | .58 (.06) | .57 (.09) | .007 | NS |
| **R Postcentral Gyrus** | ↑ | .58 (.08) | .54 (.10) | **.001** | **.01** |
| R Cuneus | ↑ | .54 (.08) | .52 (.10) | .028 | NS |
| R Occipital Gyrus | ↑ | .55 (.08) | .54 (.09) | .023 | NS |
| R Thalamus | ↑ | .54 (.08) | .51 (.10) | .017 | NS |
| **L Middle Frontal Gyrus** | ↑ | .55 (.08) | .51 (.10) | **.002** | **.02** |
| L Inferior Temporal Gyrus | ↑ | .58 (.06) | .56 (.09) | .017 | NS |
| L Fusiform Gyrus | ↑ | .59 (.07) | .58 (.09) | .007 | NS |
| L Inferior Parietal Lobe | ↑ | .59 (.06) | .57 (.08) | .036 | NS |
| L Insula | ↑ | .59 (.06) | .57 (1.0) | .016 | NS |
| *Degree* |  |  |  |  |  |
| L Post. sup. Temporal sulcus | ↑ | 15.8 (6.3) | 14.7 (6.0) | .03 | NS |
| R Postcentral Gyrus | ↑ | 17.4 (6.4) | 15.5 (6.5) | .024 | NS |
| *BC* |  |  |  |  |  |
| **R Postcentral Gyrus** | ↑ | 63.8 (42.9) | 51.5 (45.5) | **.013** | **.039** |
| L Middle Frontal Gyrus | ↑ | 55.0 (41.2) | 51.5 (64.1) | .049 | NS |
| R Insula | ↓ | 59.4 (36.7) | 67.4 (46.9) | .043 | NS |

Abbreviations: L, left; R, right; BE, Participants with Binge Eating; CON, control; p<sub>BON</sub>, bonferroni corrected p-value, Symptoms; BC, betweenness-centrality. Significant results following Bonferroni correction are shown in bold.

Compared to controls, children with BE showed significantly increased BC for successful response inhibition in the left posterior superior temporal sulcus (*F*(1) = 4.8, *p_uncorrected_ =* 0.03) and postcentral gyrus (*F*(1) = 5.2, *p_uncorrected_ =* 0.02), as well as significantly increased degree in the right postcentral gyrus (*F*(1) *= 6.3*, *p_uncorrected_ =* 0.01) and the left MFG (*F*(1) *=* 3.9, *p_uncorrected_ =* 0.05), and significantly decreased degree in the right insula (*F*(1) *=* 4.1, *p_uncorrected_ =* 0.04). Following Bonferroni correction, significance remained for the group differences in degree in the right postcentral gyrus. (**Table 2**). BMI did not significantly contribute to the ANCOVA model for degree in the right postcentral gyrus (p > 0.05).

### Correct Stop Network Hubs

Distinct patterns of hub distribution for the correct stop network were identified in the control and BE groups. According to the BC and degree measures, both groups had six common functional hubs in right SFG and right inferior frontal gyrus (IFG), right IPG, right PRG, right superior parietal gyrus (SPG) and right insula. The BE group had additional hubs in right MFG, left IPG, left insula, and left SPG, while the control group had additional hubs in right orbitofrontal gyrus, left PRG, and left fusiform gyrus. Group hubs are shown in **Figure 4**.

**Figure 4.**
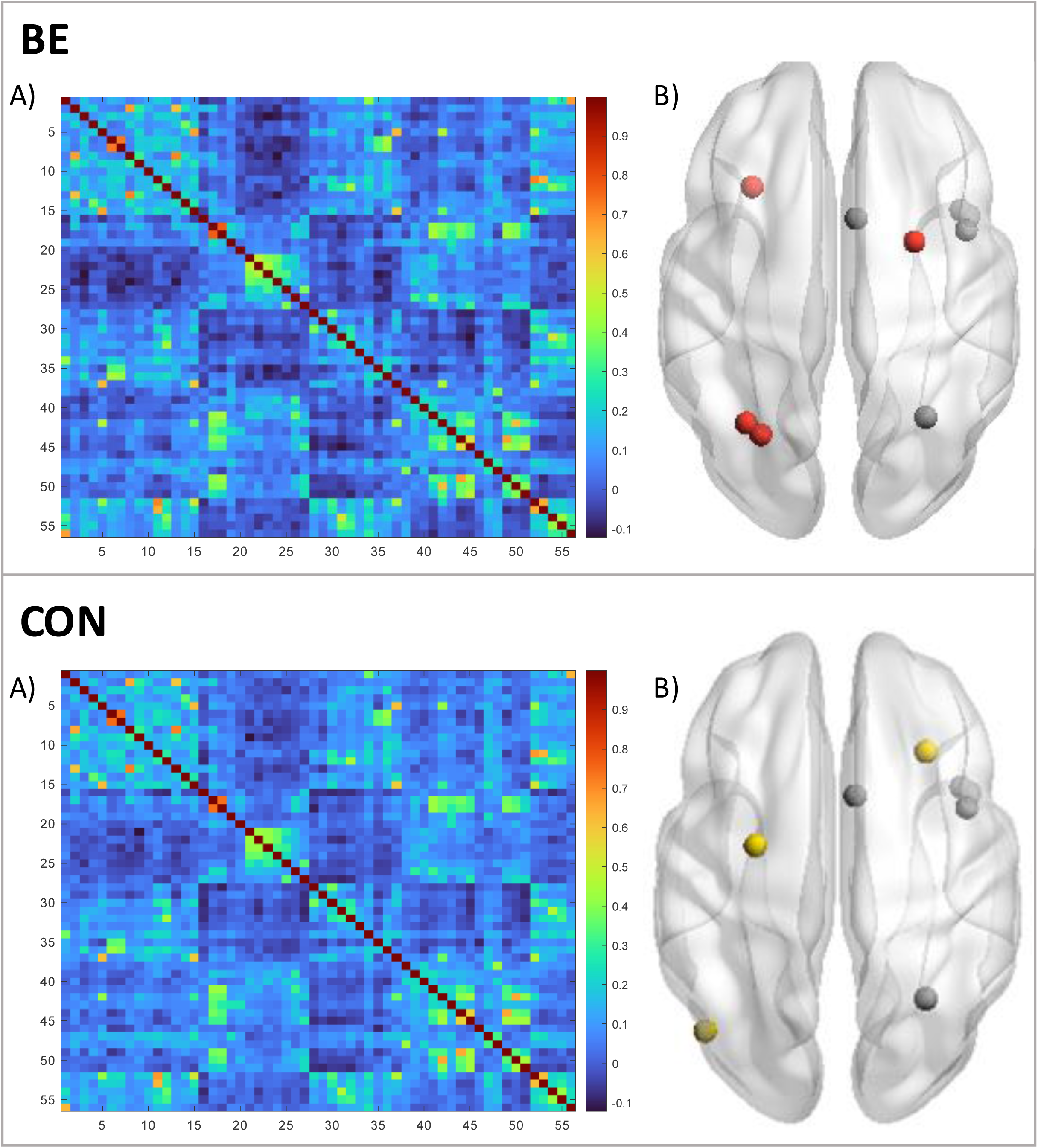
Connectivity matrices (A) and network hub nodes (B) for the correct stop network. Hub regions are defined using BC. Shared hubs are shown in grey, distinct hubs for BE and control participants are shown in red and yellow, respectively.

### Group-by-Sex Interactions in Inhibitory Control Network Nodal Properties

#### Nodal Efficiency

There was a significant group-by-sex interaction in efficiency in the left SFG (*F*(1) *=* 4.7, *p_uncorrected_ =* 0.032). Post-hoc independent samples t-test showed significantly lower efficiency in the left SFG in control girls compared to control boys (t = -2.5, p = 0.006), but no significant difference between boys and girls with BE.

#### Nodal Connectedness

Prior to Bonferroni correction, there were significant group-by-sex interactions in BC in right (*F*(1) *=* 5.9, *p_uncorrected_ =* 0.016), and left insula (*F*(1) *=* 7.5, *p_uncorrected_ =* 0.007) and in right SFG (*F*(1) *=* 5.4, *p_uncorrected_ =* 0.021) and left SPL, (*F*(1) *=* 4.1, *p_uncorrected_ =* 0.045, as well as significant group-by-sex interactions in degree in left SFG (*F*(1) *=* 6.0, *p_uncorrected_ =* 0.015) and right MFG (*F*(1) *=* 4.7, *p_uncorrected_ =* 0.021). Following Bonferroni correction, significance remained for the interactions in BC in the left insula and degree in the MFG. Post-hoc independent samples t-tests showed significantly higher BC in left insula (t = 2.8, p = 0.002) and significantly lower degree in right MFG (t = -2.2, p = 0.017) in girls with BE compared to boys with BE. In contrast, control girls had significantly lower BC in left SFG compared to control boys (t = -2.7, p = 0.004).

## DISCUSSION

This study is the first to investigate topological properties of the functional brain network for inhibitory control in preadolescent children with BE. As hypothesized, we identified several organizational in children with BE compared to control children, including significantly increased nodal efficiency of right postcentral gyrus and left middle frontal gyrus, and increased BC of right postcentral gyrus. Children with BE also showed a distinct inhibitory control network hub distribution compared to control children. Sex-by-group interactions revealed group-specific sex differences in insular and frontal cortices.

Children with BE showed increased nodal efficiency of left MFG and increased nodal efficiency and degree of right postcentral gyrus during successful inhibition of responses. Both regions have previously been highlighted as nodes of a widespread inhibitory control network (37). Specifically, MFG is suggested to be the locus of inhibitory control in humans, while increased activation in the postcentral gyrus is associated with poorer inhibition of response (37, 38). Current findings align with previous reports of increased activation in left MFG in adults with binge eating episodes (39) and in prodromal children during failed inhibitory control (17). Interestingly, these functional brain alterations were observed despite comparable behavioral performance in both cases. One interpretation of such a finding is that children with BE employ a greater level of activation in the inhibitory control network to inhibit behavioral choices.

This requirement of greater recruitment of resources to inhibit a response is reflected in the presence of additional network hubs in the IFG and insula in children with BE. An increased demand on the neurocircuitry to achieve inhibitory control may reflect a vulnerability in individuals with BE, which in the context of a highly rewarding stimulus (e.g. palatable food) may contribute to unsuccessful inhibition of eating, and the associated perception of a lack of control. Taken together, previous and current results suggest that the distinct topological organization of the inhibitory control network identified in children with BE may be a stable characteristic of individuals with a propensity to binge eat (24) Crucially, current findings in children with sub-clinical presentations of BED symptoms add to evidence of frontal cortex dysfunction during inhibition, even when a full-threshold eating disorder diagnoses is not present(17). Longitudinal research should consider the possibility of altered functional organization in the MFG and its role in the maintenance of binge eating.

We identified sex-by-group interactions in nodal efficiency and connectedness of left SFG and connectedness of left insula. Developmental sex differences in inhibitory control have previously been noted and may explain these interaction effects (20). Among adults, greater inhibitory control activation in women compared to men has been linked to increased inhibitory control over food cues (40). Given that risk for BE rises specifically for girls during adolescence, the current findings of increased BC in the left insula and decreased degree in MFG in girls versus boys with BE may reflect either specific vulnerabilities in girls or distinct protective mechanisms in boys. Interestingly, several sex differences identified in the control group were not evident in children with BE, potentially reflecting a lack of developmentally appropriate sexual dimorphism in the inhibitory control network. It is likely that any differences between boys and girls in the current cohort will increase with age, as circulating pubertal hormones drive divergence in neural development. Future research should continue to consider addressing the potential sex differences in inhibitory control within binge eating groups, particularly in longitudinal research, where increasing divergence in factors such as pubertal hormones may lead to greater changes in eating disorder prevalence, and brain network organization.

There are several limitations of this study that should be addressed. The main limitation of the current study is its cross-sectional nature. While the risk of BED increases during puberty (41), functional alterations in the inhibitory control network have been identified both in adults and children with subclinical BE and BED. Research to date, including current results, do not address the way in which organization in the inhibitory control network, and its relationship to BE, develops over the course of adolescence into adulthood. A second limitation with the current study is the difference in BMI between the BE and control groups. Inhibitory control, and its neural correlates, have been associated with BMI in adolescent participants (42, 43) and although BMI percentile was included as a covariate, and showed no significant effects in any models, the potential developmental influence of BMI on brain function should not be overlooked.

In conclusion, children with BE showed distinct topological organization of the functional network subserving inhibitory control, including increased nodal efficiency and connectedness of frontal regions. Distributional differences of the inhibitory control network hubs, defined by BC and degree, were also observed between children with BE and control children. These novel findings suggest that altered topological organization of the inhibitory control network may underlie the increased risk of the presence of binge eating episodes in children. In accordance with previous research, these systems-level functional network differences were evident despite comparable behavioral performance between the two groups, perhaps reflecting compensatory mechanisms required for successful inhibition of action in children with BE.

## Acknowledgements

This work was partially supported by NIMH (R01 MH126448). Data used in the preparation of this article were obtained from the Adolescent Brain Cognitive Development (ABCD) Study (https://abcdstudy.org), held in the NIMH Data Archive (NDA). This is a multisite, longitudinal study designed to recruit more than 10,000 children age 9-10 and follow them over 10 years into early adulthood. The ABCD Study® is supported by the National Institutes of Health and additional federal partners under award numbers U01DA041048, U01DA050989, U01DA051016, U01DA041022, U01DA051018, U01DA051037, U01DA050987, U01DA041174, U01DA041106, U01DA041117, U01DA041028, U01DA041134, U01DA050988, U01DA051039, U01DA041156, U01DA041025, U01DA041120, U01DA051038, U01DA041148, U01DA041093, U01DA041089, U24DA041123, U24DA041147. A full list of supporters is available at https://abcdstudy.org/federal-partners.html. A listing of participating sites and a complete listing of the study investigators can be found at https://abcdstudy.org/consortium_members/. ABCD consortium investigators designed and implemented the study and/or provided data but did not necessarily participate in the analysis or writing of this report. This manuscript reflects the views of the authors and may not reflect the opinions or views of the NIH or ABCD consortium investigators. The ABCD data repository grows and changes over time. The ABCD data used in this report came from 10.15154/h3rr-zp41.

