## Supplemental Material for "Altered Brain Network Topology during Successful Response Inhibition in Children with Binge Eating"

**Correct Go Network**

**Functional Network Construction**

Beta value maps were extracted for correct go trials. The total number of go trials for each subject was contingent on task performance, and the number of volumes extracted was therefore different for each participant. The average number of go trials across all participants was 241. Extracted volumes were combined sequentially to form a whole-brain beta-series, and the average beta-series values for each network node were extracted. Four-level wavelet filtering was performed on the beta-series to remove motion-related noise.

**Correct Go Network**

*Nodal Efficiency*

Group comparisons showed that, relative to control participants, participants with BE showed significantly increased nodal efficiency for response inhibition in the left middle temporal gyrus (*F*(1) *=* 4.7, *p_uncorrected_ =* .032), left middle frontal gyrus (*F*(1) = 4.4, *p_uncorrected_* = .038). These group differences did not survive Bonferroni correction.

*Nodal Connectedness*

Relative to control participants, participants with BE showed increased BC in the left nucleus accumbens (*F*(1) *=*4.3, *p_uncorrected_ =* .04), and left putamen (*F*(1) = 8.7, *p_uncorrected_ =* .004), and in the right caudate (*F*(1) *=* 4.6, *p_uncorrected_ =* .032). Following Bonferroni correction, significant group differences remained in the left putamen. Participants with BE showed significantly increased degree in the L posterior superior temporal sulcus (*F*(1) *=* 4.8, *p_uncorrected_ =* .029) and the left posterior occipital gyrus (*F*(1) = 7.6, *p_uncorrected_ =* .006). Following Bonferroni correction, significant group differences remained in the left posterior occipital gyrus.

*Correct Go Network Hubs*
Distinct hubs of the CG network were identified in the control and BE groups. Defined using degree, 12 common network hubs were identified including inferior frontal, superior frontal, and striatal nodes. Additional hubs in participants with BE were identified in the bilateral IPL, and nucleus accumbens, whereas the control group had additional hubs in the hippocampus, parahippocampal gyrus and putamen. Defined using BC, 6 common hubs between participants with BE and control subjects were identified including inferior frontal, L SFG, L MTL and R IPL. Additional hubs were identified in the BE group in the R MTL, L inferior temporal lobe, and right putamen, whereas control subjects had additional hubs in the hippocampus, parahippocampal gyrus, left putamen, right SFG and left IPL.
